# Assessments of Evoked and Spontaneous Pain Following Administration of Gabapentin and the Cannabinoid CB_2_ agonist LY2828360 in a Rat Model of Spared Nerve Injury

**DOI:** 10.1101/2025.10.29.685420

**Authors:** Kelsey G. Guenther, Jonathon D. Crystal, Andrea G. Hohmann

## Abstract

Cannabinoid CB_2_ agonists reduce stimulus-evoked behavioral hypersensitivities in preclinical pain models, but their ability to modulate spontaneous pain remains unexplored. Spontaneous pain has been assessed in rodents using the conditioned place preference (CPP) approach, given that the relief of pain is described as rewarding and results in negative reinforcement (i.e. removal of an aversive pain state). LY2828360 is a CB2 agonist that failed in a clinical trial for osteoarthritis pain. We compared impact of LY2828360 on evoked and spontaneous pain using a spared nerve injury (SNI) model in rats. First, we verified that an analgesic dose of gabapentin (100 mg/kg i.p.) produces CPP in rats with SNI, but not in sham-operated rats, consistent with a previous report (Griggs et al. 2015). We then used a within-subjects design to ascertain whether the CB_2_ agonist LY2828360 (10 mg/kg i.p., chronic) would suppress both evoked and spontaneous pain in rats with SNI. To assess evoked pain behavior, mechanical paw withdrawal thresholds were measured and revealed that LY2828360 reliably suppressed mechanical hypersensitivity in the paw *ipsilateral*, but not *contralateral* to SNI. Furthermore, efficacy was sustained across repeated injections without development of tolerance. To assess spontaneous pain behavior, we tested the ability of LY2828360 to prevent gabapentin-induced CPP in the SNI model, as failure to develop CPP to gabapentin following treatment with an analgesic has been considered evidence of suppression of spontaneous pain. The same rats that showed suppression of mechanically-evoked responses following chronic LY282860 treatment did not develop CPP to gabapentin. However, rats that were tested in parallel and treated chronically with vehicle showed robust mechanical hypersensitivity, but also did not develop CPP to gabapentin. These studies document that CB_2_ agonist-induced suppression of mechanically evoked pain is highly robust and reproducible, whereas CPP, used to assess spontaneous pain, is vulnerable to disruption and requires rigorous controls to rule out alternative explanations (e.g. failure to learn).

## 1. Introduction

Pain is a complex disorder involving sensory, cognitive, and affective-motivational components. The affective component of pain is reported as the most distressing to patients, but most pre-clinical studies measure only somatosensation. Specifically, most preclinical studies employing rodent pain models measure withdrawal responses to an applied sensory stimulus (evoked pain), whereas most patients with neuropathic pain instead report spontaneous pain (ongoing) which motivates the consumption of analgesic compounds and can induce negative affect (Rice et al., 2018). Lack of research regarding the spontaneous component of pain is primarily due to the difficulty of assessing this pain symptom in animal models (Navratilova et al., 2013). Operant paradigms may provide alternative experimental methods to assess spontaneous pain while also providing higher translatability than traditional pain assessment methods, particularly through the measurement of pain-motivated behavior in animals. It is possible to assess an animal’s motivation to avoid or reduce pain using behavioral methods such as conditioned place preference (CPP) or drug self-administration which can determine the ability of analgesic compounds to be negatively reinforcing (Cabañero et al., 2020; Gutierrez et al., 2011; Gutierrez et al., 2021; King et al., 2009; Martin & Ewan, 2008).Reliance on CPP in rodent pain models is complicated by the possibility that absence of CPP may reflect the ineffectiveness of a putative pain therapeutic or the failure to learn the chamber association. Our work suggests that a combination of assessments of spontaneous pain, evoked pain, and CPP provides a powerful combination for drawing valid conclusions about putative pain therapeutics.

CB_2_ receptor activation has been shown to suppress evoked pain (behavioral hypersensitivity evoked in response to a mechanical stimulus) in pre-clinical models (Cabañero et al., 2021; Guindon & Hohmann, 2008), but its ability to modulate spontaneous pain has not been fully explored. Our lab previously showed that rats with SNI-induced neuropathy will self-medicate with the CB_2_ receptor agonist AM1241, an effect that is dependent on the CB_2_ receptor (Gutierrez et al., 2011). Another group showed that mice with partial sciatic nerve ligation (PSNL) will self-medicate with the CB_2_ receptor agonist JWH-133 in an intravenous self-administration paradigm (Cabañero et al., 2020). To our knowledge, ability of CB_2_ receptor agonists to produce preference in the CPP paradigm in animals with neuropathic pain has not been investigated. The CB_2_ receptor agonist LY2828360 shows antinociceptive efficacy in rodent models of neuropathic pain (Carey et al., 2023; Guenther et al., 2024; Lin et al., 2018; Lin et al., 2022) and reduces pain behavior in a spared nerve injury (SNI) model in rats (Guenther et al., 2025). Additionally, LY2828360 itself does not produce drug-context association, indicative of reward, in the CPP paradigm in naïve rats (Guenther et al., 2025). While it is clear LY2828360 can mitigate evoked pain behavior, the ability of this compound to modulate spontaneous pain behavior has not been assessed. These investigations are important because LY2828360 failed for efficacy in a Phase 2 clinical trial of osteoarthritis pain (Pereira et al., 2013).

Here, we use a within subject design to test the ability of LY2828360 to reduce both evoked and spontaneous pain behaviors in the same rats with SNI. Given that repeated administration of LY2828360 produces a sustained antinociceptive effect up to 24 hours after the previous injection (Guenther et al., 2025), we examined the ability of chronic LY28282360 to prevent the development of CPP to the shorter acting analgesic compound gabapentin, instead of testing LY2828360 alone in this paradigm, given the necessity of daily drug administration in this paradigm. Gabapentin is a first line treatment for human patients suffering from neuropathic pain (Attal et al., 2010) and can induce CPP in various pre-clinical neuropathic pain models (Griggs et al., 2015; Park et al., 2013; Wagner et al., 2014). Therefore, we first determined whether we could replicate the findings of (Griggs et al., 2015) to show that gabapentin (100 mg/kg) produces CPP in rats with SNI-induced neuropathy. Then, we evaluated the ability of an analgesic dose of LY2828360 (10 mg/kg i.p., chronic) to prevent this gabapentin-induced CPP under conditions in which rats were chronically pre-treated with either vehicle of LY2828360. Importantly, responses to evoked pain were assessed in the same animals. Our studies document that CPP, which depends on learning, can be highly sensitive to disruption even when reference experiments are replicated.

## 2. Methods

### 2.1. Subjects

Adult male Sprague-Dawley rats (Envigo, Indianapolis, IN) between the age of 90-100 days at the start of experiments were used. Rats were single-housed in a temperature-controlled colony room on a 12/12-hr light/dark cycle (lights on at 8am; lights off at 8pm). All behavioral testing occurred during the dark cycle under red light. Rats were maintained on ad libitum food and water. A single experimenter (KG) performed all experimental manipulations including daily handling, surgical procedures, CPP studies and assessment of mechanical paw withdrawal thresholds. All experiments were approved by the Indiana University Animal Care and Use Committee.

### 2.2. Drugs

LY2828360 (8-(2-chlorophenyl)-2-methyl-6-(4-methylpiperazin-1yl)-9-(tetrahydro-2H-pyran-4-yl)-9H-purine) was synthesized by Sai Biotech (Mumbai, India; purity >98%). LY2828360 was dissolved in a vehicle consisting of 3% dimethylsulfoxide (DMSO; Sigma-Aldrich, St. Louis, MO), and the remaining 97% consisted of emulphor (Alkamuls EL-620; Solvay), 95% ethanol (Sigma-Aldrich) and 0.9% saline (Aqualite System; Hospira, Inc., Lake Forest, IL) at a 1:1:18 ratio for a 10 mg/kg intraperitoneal (i.p.) injection at a volume of 2 ml/kg (5 mg/ml concentration). Gabapentin was dissolved in saline for a 100 mg/kg i.p. injection at a volume of 1 ml/kg (100 mg/ml concentration).

### 2.3. Spared nerve injury surgery

SNI surgery was performed in rats to induce neuropathic nociception in the ipsilateral paw as described previously (Decosterd & Woolf, 2000; Gutierrez et al., 2011; Gutierrez et al., 2021). Briefly, rats were deeply anesthetized with isoflurane, the flank of the rat was shaved and prepared for aseptic surgery. Then, a small incision (approximately 1.5 cm) was made on the flank of the rat and the underlying biceps femoris muscle was gently separated to expose the sciatic nerve. The common peroneal and tibial branches of the sciatic nerve were tightly ligated with a silk suture (5-0 PERMA-HAND Silk Suture, Ethicon) then cut, leaving the sural branch intact. The muscle was then closed and held together with a single silk suture before closing the skin incision with nylon sutures (4-0 ETHILON Black Monofilament, Ethicon), which were then removed 1-week post-surgery. Sham surgeries were also performed and involved separating of the muscle to expose the sciatic nerve, but no ligation or cutting of the nerve occurred. Rats were not tested during the initial 2-week period following SNI surgery to allow them to recover from surgery and the acute inflammatory phase of SNI. Baseline mechanical paw withdrawal thresholds were measured before and after the induction of neuropathic nociception to ensure the development of mechanical hypersensitivity.

### 2.4. Assessment of mechanical hypersensitivity

Mechanical paw withdrawal thresholds were measured using an electronic von Frey anesthesiometer (IITC model Alemo 2290-4, Woodland Hills, CA) as described in our previously published work (Gutierrez et al. 2011; Gutierrez et al. 2021). Rats were placed on a testing table with a stainless-steel wire mesh flooring (1.2 x 1.2 cm gaps in wire mesh) and confined under a clear Plexiglas chamber (13.5 x 13.5 x 23.5 cm) for at least 30 minutes before testing. Following habituation, a force was applied with the von Frey analgesiometer to the hind paw until a paw withdrawal response was observed. The force was applied toward the lateral edge of the paw to stimulate the site of injury of the SNI model (Cichon et al., 2018). Each paw was measured twice at each time point in a right-left-right-left order with an interstimulation interval of several minutes between paws. Specifically, the right paw was tested for all rats before moving on to test the left paw for all rats and so on for each time point; testing the entire right-left-right left sequence for any given rat took approximately 15 minutes. Paw withdrawal thresholds (measured in duplicate) were averaged separately for the paw ipsilateral and contralateral to SNI.

### 2.5. Conditioned place apparatus

A custom-configured three chamber apparatus was used for CPP (Med-Associates Inc., Fairfax, VT, USA) to permit assessment of place preference using an unbiased approach. Two conditioning chambers (21 x 21 x 28 cm) on either side were separated via guillotine doors from a central neutral (gray) chamber (12 x 21 x 21 cm). The conditioning chambers had distinct visual cues (thick vertical or thin horizontal black and white stripes) and were custom configured to have an equivalent amount of black and white in both chambers to eliminate chamber bias. The time spent in each chamber was automatically recorded via Med-Associates software over a 15-minute interval during habituation, pre-test and CPP test days and over a 30-minute interval during all conditioning days.

### 2.6. Experiment 1: Gabapentin-induced place preference in the SNI model

The experimental protocol for Experiment 1 is shown in **Figure 1A**. Mechanical paw withdrawal thresholds in Experiment 1 were measured 1 day prior to SNI surgery and 2 weeks after SNI surgery (i.e., prior to the start of CPP). Following completion of the CPP protocol, paw withdrawal thresholds were measured 24 h and 72 h later in a crossover design to assess the time course of analgesic efficacy of gabapentin vs vehicle that was performed. In the crossover experiment, half the rats received gabapentin (100 mg/kg i.p.) and the other half received saline on the first day. Two days later, each group received the opposite i.p. treatment. Paw withdrawal thresholds were measured at 0.5, 1, 2 and 4 hours after each injection.

**Fig. 1.**
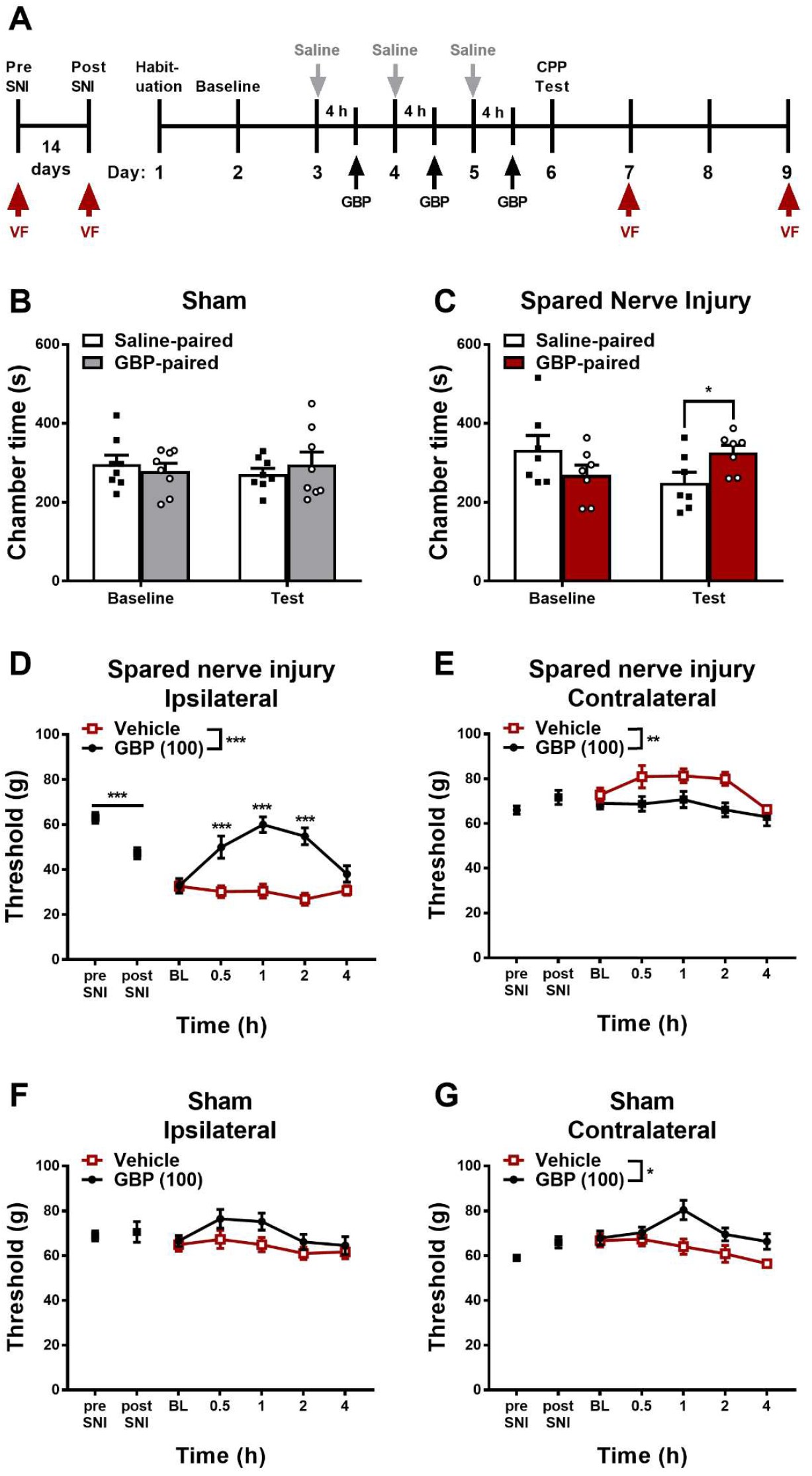
Gabapentin (GBP) induced conditioned place preference (CPP) and suppressed mechanical hypersensitivity in male rats with SNI-induced neuropathy. Schematic shows the timeline of the von Frey (VF) measurements and conditioned place preference study (**A**). Gabapentin did not produce place preference or aversion relative to a saline chamber in sham operated rats (**B**), but did induce CPP in male rats with SNI-induced neuropathy (**C**). Gabapentin (100 mg/kg i.p.) increased mechanical paw withdrawal thresholds in the paw *ipsilateral* (**D**) and *contralateral* (**E**) to SNI. Gabapentin increased mechanical paw withdrawal thresholds in the paw *contralateral* (**G**) but not *ipsilateral* (**F**) to sham surgery. n = 7-8 per group. Data expressed as Mean ± SEM. Two-way ANOVA followed by one-tailed paired t-test (CPP) or Sidak’s multiple comparison *post-hoc* test (mechanical paw withdrawal thresholds). \**p* < 0.05, \*\**p* < 0.005, \*\*\**p* < 0.001.

The CPP protocol was initiated 14 days after SNI surgery. On day 1 of the CPP experiment, rats were placed in the middle chamber and allowed to freely roam between all three chambers to habituate to the apparatus. On day 2, rats were again allowed to freely roam the three chambers for 15 minutes after which baseline preference assessment was conducted to confirm that rats did not show a bias for either chamber prior to drug pairing. Rats who spent more than 720 seconds (i.e. 80%) or less than 180 seconds (i.e. 20%) of the baseline trial in either chamber were excluded from the study. The remaining rats were randomly assigned to receive active treatments in the left or right chamber (Sham: n = 8; SNI n = 7). Next, rats underwent three days of conditioning trials during which they received saline (i.p.) and then were immediately restricted to a single chamber of the CPP apparatus for 30 minutes in the morning. Then, 4 hours after the saline injection, rats received gabapentin (100 mg/kg i.p.) before being restricted to the opposite chamber of the CPP apparatus for 30 minutes. On day 6, rats were placed in the middle chamber with guillotine doors open and allowed to freely roam between all three chambers to assess the impact of treatment on CPP in a drug-free state for 15 minutes.

### 2.7. Experiment 2: Impact of LY2828360 on gabapentin-induced place preference in the SNI model

In Experiment 2 all rats were subjected to surgical procedures to induce SNI. Mechanical paw withdrawal thresholds were measured 1 day prior to SNI surgery as well as 2 weeks after SNI surgery (i.e., prior to the start of CPP and chronic drug treatment). The CPP protocol was initiated 14 days after SNI surgery. On day 1, rats were placed in the middle chamber and allowed to freely roam between all three chambers to habituate to the apparatus. On day 2, rats were again allowed to freely roam the three chambers for 15 minutes after which baseline preference assessment was conducted to confirm that rats did not show a bias for either chamber prior to drug pairing. On day 3 rats began chronic dosing with either vehicle (n = 11) or LY2828360 (10 mg/kg) (n = 10) in which they received once daily i.p. injections. This dose was selected based upon our previous work showing that repeated administration of LY2828360 (10 mg/kg i.p.) reduces mechanical hypersensitivity in rats with SNI (Guenther et al., 2025). On day 6 (i.e., day 4 of chronic dosing), mechanical paw withdrawal thresholds were measured before (pre-drug baseline) and 1-hour post-injection of vehicle or LY2828360. On day 7, the three-day conditioning phase began. The chamber pairings on conditioning days were identical to those in Experiment 1, with the exception that rats received vehicle or LY2828360 (10 mg/kg i.p.) 1 hour prior to the morning (saline-paired) conditioning trial. On the test day, no injection of vehicle or LY2828360 occurred, and rats were allowed to freely roam between all three chambers to assess the impact of treatment on CPP in a drug-free state for 15 minutes. Paw withdrawal thresholds were additionally measured 24 h after the CPP test day. Paw withdrawal measurements were taken to confirm that SNI-induced mechanical hypersensitivity developed and document that LY2828360 (10 mg/kg i.p. chronic) treatment suppressed mechanical hypersensitivity in the same rats used in the CPP study.

### 2.8. Statistical Analysis

Data were analyzed by Two-way repeated measures ANOVA followed by one-tailed paired t-tests for CPP data and Sidak’s multiple comparisons post-hoc test for mechanical paw withdrawal thresholds to determine significant differences between groups. Preference scores for CPP data were calculated (test time – baseline time) and analyzed by one-tailed paired t-tests. Statistical analysis was performed using Graphpad prism 7 (Graphpad prism software, San Diego, CA).

## 3. Results

### 3.1. Experiment 1: Gabapentin-induced place preference in the SNI model

In the CPP paradigm, sham-operated rats did not exhibit differences in chamber preference time irrespective of drug pairing or conditioning phase and the interaction was not significant (Drug pairing: F_1,14_ = 0.01962, *p* = 0.8906; Conditioning phase: F_1,14_ = 0.04174, *p* = 0.8410; Interaction: F_1,14_ = 0.8898, *p* = 0.3615) (**Fig. 1B**). In rats with SNI, a significant interaction was observed after conditioning whereas drug pairing and conditioning phase did not alter chamber preference time overall (Drug pairing: F_1,12_ = 0.03821, *p* = 0.8483; Conditioning phase: F_1,12_ = 0.3838, *p* = 0.5472; Interaction: F_1,12_ = 9.992, *p* = 0.0082) (**Fig. 1C**). Two-tailed paired t-test showed no difference in time spent in either chamber prior to conditioning in rats with SNI (*p* = 0.2728). By contrast, on the test day (following conditioning), rats with SNI spent more time in the gabapentin-paired chamber compared to the saline-paired chamber (*p* = 0.0454) (**Fig. 1C**). Analysis of preference scores similarly confirmed that rats with SNI-induced neuropathy showed a preference for the gabapentin-paired chamber (*p* = 0.0313; one-tailed t-test), but rats that received sham surgery did not (*p* = 0.1954, one-tailed t-test) (**Fig. 2**).

**Fig. 2.**
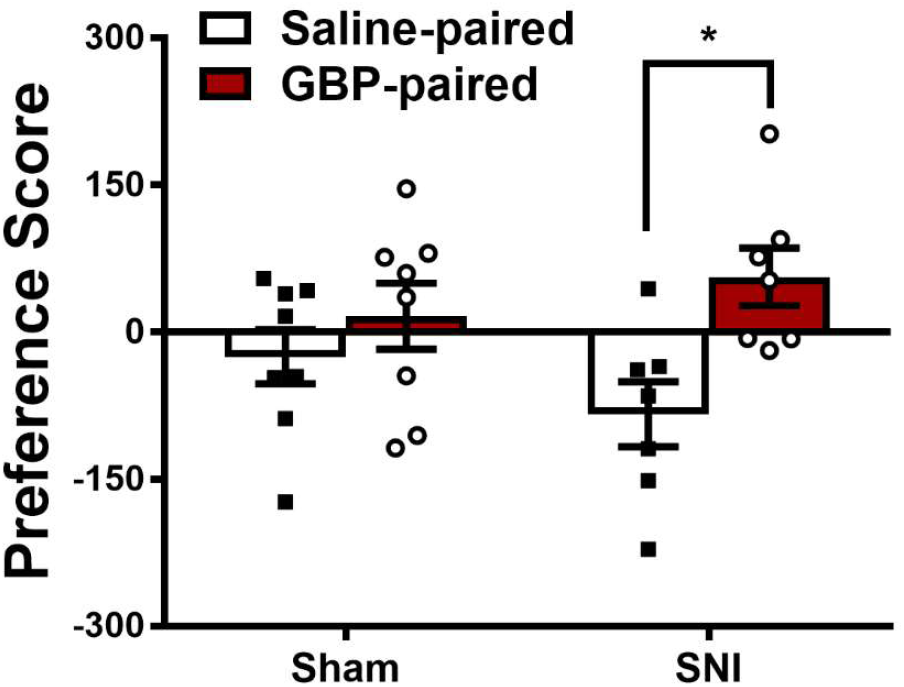
Gabapentin (GBP; 100 mg/kg i.p.) increased preference scores in male rats with SNI-induced neuropathy but not in sham-operated rats. Preference scores were calculated as post-conditioning time – pre-conditioning time in the drug paired chamber. Data are expressed as Mean ± SEM. One-tailed paired t-test. \**p* < 0.05.

Prior to pharmacological treatment, paw withdrawal thresholds were lowered in the *ipsilateral* paw of rats with SNI relative to pre-surgery baseline (*p* = 0.0008) (**Fig. 1D**), but no difference was observed in the *contralateral* paw (*p* = 0.1576) (**Fig. 1E**). In sham-operated rats, prior to pharmacological treatment paw withdrawal thresholds were unchanged in the *ipsilateral* (*p* = 0.6722) (**Fig. 1F**) and *contralateral* paw relative to pre-surgery baseline (*p* = 0.0852) (**Fig. 1G**).

In the paw *ipsilateral* to SNI, gabapentin treatment (100 mg/kg i.p.) increased mechanical paw withdrawal thresholds relative to vehicle, paw withdrawal thresholds changed across time and the interaction was significant (Group: F_1,16_ = 37.32, *p* < 0.0001; Time: F_4,64_ = 6.242, *p* = 0.0003; Interaction: F_4,64_ = 9.995, *p* < 0.0001). Sidak’s multiple comparisons *post hoc* test revealed that gabapentin increased mechanical paw withdrawal thresholds in the paw *ipsilateral* to SNI at 0.5 (*p* = 0.0002), 1 (*p* < 0.0001) and 2 hours (*p* < 0.0001) post-injection (**Fig. 1D**). In the paw *contralateral* to SNI, gabapentin treatment (100 mg/kg i.p.) also increased mechanical paw withdrawal thresholds relative to vehicle, paw withdrawal thresholds changed across time and the interaction was not significant (Group: F_1,16_ = 16.11, *p* = 0.0010; Time: F_4,64_ = 3.719, *p* = 0.0088; Interaction: F_4,64_ = 1.089, *p* < 0.3695) (**Fig. 1E**). In the paw *ipsilateral* to sham surgery, gabapentin treatment (100 mg/kg i.p.) did not reliably increase mechanical paw withdrawal thresholds relative to vehicle, paw withdrawal thresholds changed across time and the interaction was not significant (Group: F_1,16_ = 3.648, *p* = 0.0742; Time: F_4,64_ = 3.481, *p* = 0.0123; Interaction: F_4,64_ = 0.8179, *p* = 0.5185) (**Fig. 1F**). In the paw *contralateral* to sham surgery, gabapentin treatment (100 mg/kg i.p.) increased mechanical paw withdrawal thresholds relative to vehicle, paw withdrawal thresholds changed across time and the interaction was not significant (Group: F_1,16_ = 7.985, *p* = 0.0122; Time: F_4,64_ = 4.141, *p* = 0.0048; Interaction: F_4,64_ = 2.332, *p* = 0.0652) (**Fig. 1G**).

### 3.2. Experiment 2: Impact of LY2828360 on gabapentin-induced place preference in the SNI model

The experimental protocol for Experiment 2 is shown in **Figure 3A**. Note that prior to pharmacological treatment, paw withdrawal thresholds were lowered in the *ipsilateral* paw of rats with SNI relative to pre-surgery baseline (*p* < 0.0001) (**Fig. 3B**). By contrast, paw withdrawal thresholds were modestly increased in the paw *contralateral* to SNI (*p* = 0.0019) (**Fig. 3C**), consistent with changes in weight bearing.

**Fig. 3.**
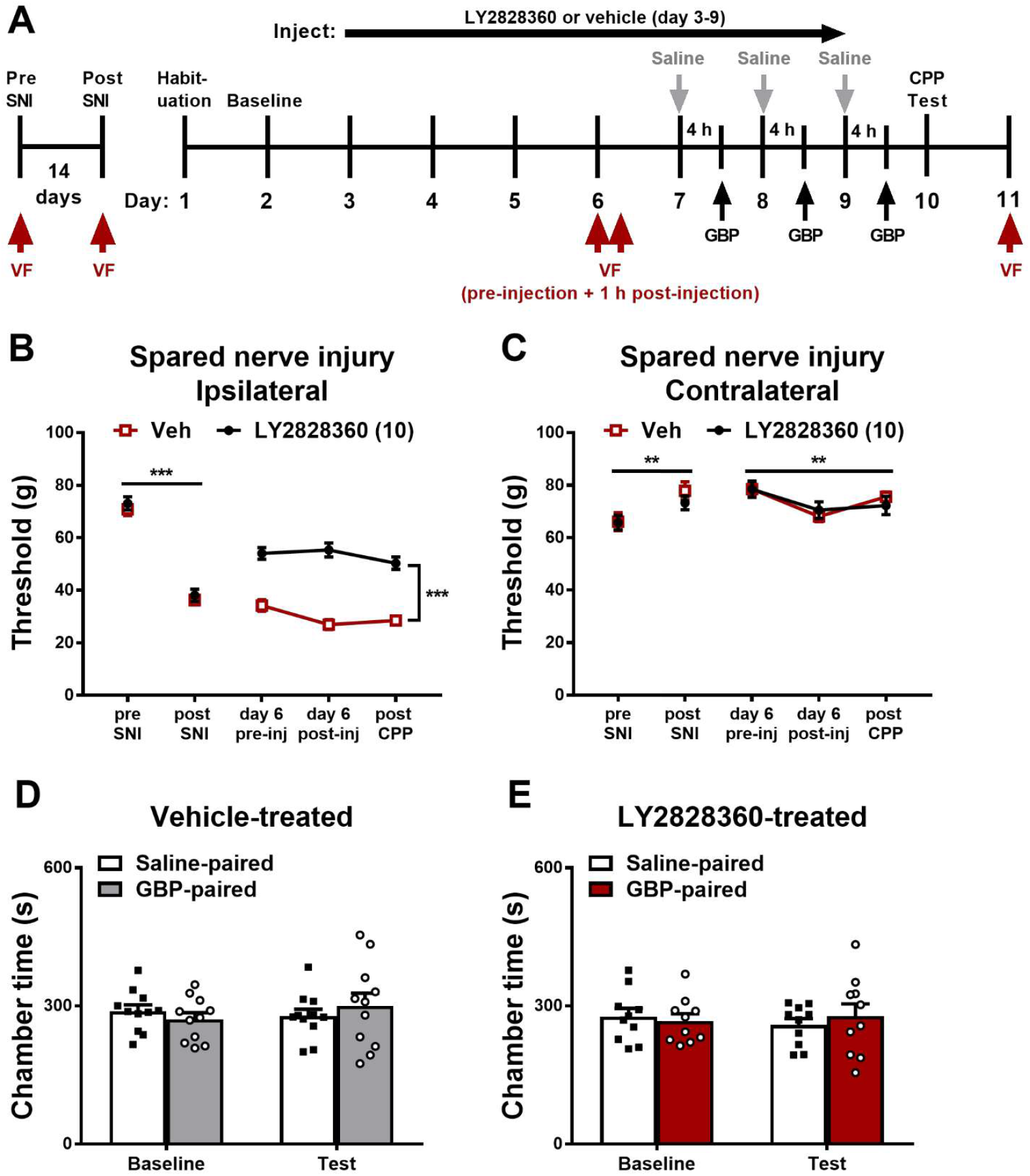
Repeated administration of LY2828360 suppressed mechanical hypersensitivity relative to vehicle in male rats with SNI, and gabapentin (GBP) did not produce conditioned place preference (CPP) in SNI rats that were pre-treated with either vehicle or LY2828360. Schematic shows the timeline of pharmacological manipulations, von Frey (VF) measurements and conditioned place preference study (**A**). LY2828360 (10 mg/kg i.p.) increased mechanical paw withdrawal thresholds in the paw *ipsilateral* (**B**) but not *contralateral* (**C**) to SNI. Gabapentin (100 mg/kg i.p.) did not produce place preference or aversion relative to a saline chamber in rats with SNI-induced neuropathy when pre-treated with vehicle (**D**) or LY2828360 (10 mg/kg i.p.) (**E**). n = 10-11 per group. Data expressed as Mean ± SEM. Two-way ANOVA followed by Sidak’s multiple comparison *post-hoc* test (mechanical paw withdrawal thresholds). \*\**p* < 0.005, \*\*\**p* < 0.001.

In the paw *ipsilateral* to SNI, repeated treatment with LY2828360 (10 mg/kg i.p.) increased mechanical paw withdrawal thresholds relative to vehicle, paw withdrawal thresholds did not change across time and the interaction was not significant (Group: F_1,23_ = 135.5, *p* < 0.0001; Time: F_2,64_ = 2.893, *p* = 0.0655; Interaction: F_2,64_ = 2.561, *p* = 0.0882) (**Fig. 3B**). In the paw *contralateral* to SNI, repeated treatment with LY2828360 (10 mg/kg i.p.) did not increase mechanical paw withdrawal thresholds relative to vehicle, paw withdrawal thresholds changed across time and the interaction was not significant (Group: F_1,23_ = 0.02145, *p* = 0.8848; Time: F_2,64_ = 6.173, *p* = 0.0042; Interaction: F_2,64_ = 0.6203, *p* = 0.5422) (**Fig. 3C**). Thus, LY2828360 reliably suppressed evoked pain behavior in the injured paw, consistent with suppression of mechanical allodynia.

In the CPP test, no difference in time spent in the saline-vs. gabapentin-paired chamber was observed in a rat model of SNI-induced neuropathy when rats were pre-treated with either vehicle (Drug pairing: F_1,40_ = 0.01583, *p* = 0.9005; Conditioning Phase: F_1,40_ = 0.2203, *p* = 0.6413; Interaction: F_1,40_ = 1.124, *p* = 0.2955) (**Fig. 3D**) or LY2828360 (Drug pairing: F_1,36_ = 0.06067, *p* = 0.8068; Conditioning phase: F_1,36_ = 0.04195, *p* = 0.8389; Interaction: F_1,36_ = 0.5407, *p* = 0.4669) (**Fig. 3E**). Analysis of preference scores showed no preference for the gabapentin-paired chamber in either the vehicle-treated (*p* = 0.1218; one-tailed t-test) or LY2828360-treated (*p* = 0.4663, one-tailed t-test) groups (**Fig. 4**).

**Fig. 4.**
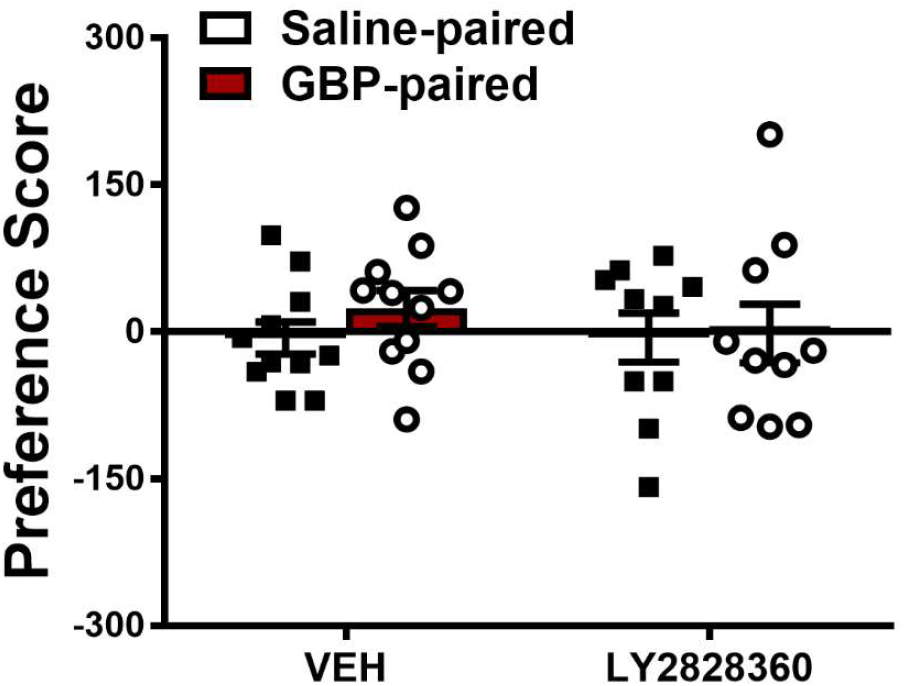
Preference scores for the gabapentin (GBP; 100 mg/kg i.p.) and saline paired chamber did not differ in SNI rats treated with either vehicle of LY2828360 (10 mg/kg i.p.). Preference scores were calculated as post-conditioning time – pre-conditioning time in the drug paired chamber. Data are expressed as Mean ± SEM. One-tailed paired t-test.

## 4. Discussion

Our studies aimed to compare effects of LY2828360, a CB2 agonist which failed for efficacy in a Phase 2 clinical trial on evoked and spontaneous pain in a rat model of spared nerve injury. The present studies document that the CB2 agonist LY2828360 produces a robust suppression of evoked pain behavior in rats with SNI, and these effects were restricted to the ipsilateral (injured) paw. We recently reported that repeated administration of LY2828360 (10 mg/kg i.p.) produces anti-allodynic efficacy that can last up to 24 hours after the prior injection in male rats (Guenther et al., 2025). These observations are robust and replicated in the current study (i.e., in which rats were treated (i.p.) with LY2828360 or vehicle for 7 consecutive days) which was designed to permit assessment of evoked and spontaneous pain in the same subjects. It was not viable to evaluate LY2828360 on its own (i.e. analogous to assessing CPP to gabapentin with repeated drug-chamber pairings, given the requirement for repeated drug dosing across conditioning trials. Therefore, we instead determined whether the anti-allodynic effects of LY2828360 following repeated once daily injections would prevent gabapentin-induced CPP in rats with SNI-induced neuropathy. However, we observed that when rats were repeatedly pre-treated with either vehicle or LY2828360, we did not observe the phenomenon of gabapentin-induced CPP. This result contrasts with our gabapentin replication study, as we expected to see gabapentin-induced CPP in the vehicle condition using our unbiased paradigm comparable to what was reported previously using a biased CPP protocol (Griggs et al., 2015). Given the failure of our positive control to produce CPP, it is not possible to conclude that LY2828360 either blocked (or failed to block) spontaneous pain. Rather, we conclude that the absence of CPP may reflect a failure to learn the chamber association in the chronic dosing groups. This interpretation must be considered in any study using CPP to assess spontaneous pain.

Several differences in the experimental methods between the two studies may have contributed to the lack of gabapentin-induced CPP seen in SNI rats receiving repeated vehicle (i.p.) treatments. First, our studies used an unbiased CPP approach. Second, methodological differences exist between the studies: there was a four-day gap between the baseline trial and the conditioning trial (i.e., to accommodate the chronic vehicle and LY2828360 injections) and additional i.p. injections occurred every morning an hour prior to chamber pairings (i.e. saline i.p. injection/saline chamber paring) which did not occur in Experiment 1. Both factors could have contributed to a failure to learn. Additionally, in the gabapentin control study (Experiment 1), conditioning began approximately two weeks post-surgery (17 days), whereas in the LY2828360 or vehicle pre-treatment study (Experiment 2) conditioning began three weeks post-surgery. Previous work has suggested that pregabalin, an anticonvulsant with a mechanism similar to gabapentin, produced CPP in the SNL model when conditioning trials started two weeks post-surgery, but CPP did not develop if conditioning began four weeks post-surgery (Asaoka et al., 2018). It is possible that the later post-surgical start time in the LY2828360 experiment (Experiment 2) contributed to the failure to establish CPP in the vehicle treated group. However, this latter hypothesis runs counter to a recent review of the SNI model, which suggest that negative affective states only develop several weeks post SNI surgery (Guida et al., 2020). It is also possible that repeated mechanical/cold stimulation is required to sensitize animals to permit assessments of ongoing (spontaneous) pain. Nonetheless, we were able to document that rats used in the CPP studies exhibited SNI-induced mechanical hypersensitivity, which was reliably suppressed by LY2828360. Moreover, LY2828360-induced suppression of SNI-induced mechanical hypersensitivity also persisted 24 hours following the CPP test (48 hours after the last injection). These studies suggest that CB_2_ agonist-induced suppression of mechanically-evoked pain is highly robust, and reproducible whereas the unbiased CPP paradigm used here to assess spontaneous pain, is inconsistent across studies and vulnerable to slight changes in experimental protocol. Additionally, our results highlight the importance of including both positive and negative controls and carefully matching experimental protocols to ensure that simpler, alternative explanations (e.g. failure to learn) can be excluded.

An important feature of our CPP experiments is that we used an unbiased approach wherein one drug-pairing chamber contained black and white vertical stripes, the other drug-pairing chamber consisted of black and white horizontal stripes separated by a central (gray) compartment. Thus, in our study each drug-pairing chamber has equivalent amounts of black and white, with different orientations. This design is important given that rats innately prefer black compared to white chambers. We largely replicate the findings obtained with a biased CPP approach (Griggs et al., 2015) showing that gabapentin (100 mg/kg i.p.) produces CPP selectively in rats with SNI, presumably by attenuating spontaneous pain associated with traumatic nerve injury using our unbiased approach. This latter study used a biased assignment approach, in addition to employing different olfactory cues (i.e. different scented chapsticks) and different intensities of lighting in a CPP apparatus containing black and white drug-pairing chambers. Specifically, rats with SNI-induced neuropathy spent more time in a gabapentin-paired chamber compared to a saline-paired chamber, whereas sham-operated rats did not show any preference for either chamber in both studies. Several other studies have reported pain relief to be negatively reinforcing by producing drug-context associations in the CPP paradigm. Clonidine and -conotoxin administered into the spinal cord and lidocaine administered in the rostral ventromedial medulla produce CPP in the spinal nerve ligation (SNL) model (King et al., 2009). However, one limitation of single trial CPP studies administering intrathecal agents such as lidocaine to induce CPP, is that the approach includes an unavoidable temporal confound; the first (e.g., morning) intrathecal pairing is always with vehicle and the second (e.g., afternoon) chamber pairing is always with lidocaine (or the drug that induces the CPP). Gabapentin is also reported to produce CPP in mice with cisplatin-induced neuropathic pain (Park et al., 2013). Nerve block with lidocaine or bupivacaine produces CPP in rats with a hind paw incision (Dalm et al., 2015; Navratilova et al., 2013) and can activate ventral tegmental area (VTA) dopaminergic cells and increase dopamine release in the nucleus accumbens (NAc) in this test (Navratilova et al., 2013). Gabapentin also produced dopamine release in the NAc in rats with SNL, and the development of gabapentin-induced CPP is blocked with inhibition of the endogenous opioid system in the anterior cingulate cortex (ACC), an area involved in pain perception (Bannister et al., 2017). Lastly, inhibition of excitatory signaling in the ACC produces CPP in the complete Freud’s adjuvant (CFA) model of persistent inflammatory pain (Kang et al., 2017). The use of the CPP paradigm to examine the motivational component of different analgesic drugs is, consequently, a promising direction for inferring the presence of spontaneous pain in pre-clinical models. In this realm, the spinal nerve ligation (SNL) model may provide a more robust phenomenon of spontaneous pain compared to the surgically sterile partial axotomy of the spared nerve injury (SNI) model.

We conclude that the present studies provide no evidence either for or against a role for LY2828360 in suppressing spontaneous pain in a CPP paradigm because in the positive control group, gabapentin did not produce CPP in rats with SNI that were treated daily with vehicle in lieu of LY2828360. Without a matched vehicle i.p. repeated dosing group, one could erroneously conclude that LY28282360 pretreatment was sufficient to block spontaneous pain such that gabapentin-induced CPP was not observed in rats with SNI under the conditions of LY2828360 pre-treatment. Moreover, these studies demonstrate the instability of the CPP paradigm for investigating spontaneous pain compared to measurements of evoked pain such as the von Frey test which are shown here to be highly replicable and reliable. Although it is important to investigate spontaneous pain in pre-clinical models given its relevance for translatability to human patients, the potential vulnerability of CPP to experimental protocol must be considered. More work should be performed using CPP methods to better understand the therapeutic potential of analgesic compounds, potentially using direct intracranial or intrathecal injections that permit faster onset of analgesic action, factors which can be expected to facilitate learning associations between behavioral context and drug treatment. Furthermore, because the mechanisms involved in non-evoked pain may differ from those of evoked pain it is important to gain an in depth understanding of the rewarding effect of pain relief and limitations of existing experimental approaches, to further the goal of developing and validating more effective analgesic strategies.

## Acknowledgments

Supported by DA047858 and DA009158 (to AGH) and an Indiana Addiction Grand Challenge Grant (to AGH and JDC). KG was supported by the Harlan Research Scholars program.

## Notes

### Competing Interest Statement

The authors have declared no competing interest.

## References

Asaoka, Y., Kato, T., Ide, S., Amano, T., & Minami, M. (2018). Pregabalin induces conditioned place preference in the rat during the early, but not late, stage of neuropathic pain. Neuroscience Letters, 668, 133–137. 10.1016/j.neulet.2018.01.029

Attal, N., Cruccu, G., Baron, R., Haanpää, M., Hansson, P., Jensen, T. S., & Nurmikko, T. (2010). EFNS guidelines on the pharmacological treatment of neuropathic pain: 2010 revision. European Journal of Neurology, 17(9). 10.1111/j.1468-1331.2010.02999.x

Bannister, K., Qu, C., Navratilova, E., Oyarzo, J., Xie, J. Y., King, T., Dickenson, A. H., & Porreca, F. (2017, Dec). Multiple sites and actions of gabapentin-induced relief of ongoing experimental neuropathic pain. Pain, 158(12), 2386–2395. 10.1097/j.pain.0000000000001040

Cabañero, D., Martin-Garcia, E., & Maldonado, R. (2021, Aug). The CB2 cannabinoid receptor as a therapeutic target in the central nervous system. Expert Opin Ther Targets, 25(8), 659–676. 10.1080/14728222.2021.1971196

Cabañero, D., Ramirez-Lopez, A., Drews, E., Schmole, A., Otte, D. M., Wawrzczak-Bargiela, A., Huerga Encabo, H., Kummer, S., Ferrer-Montiel, A., Przewlocki, R., Zimmer, A., & Maldonado, R. (2020, Jul 20). Protective role of neuronal and lymphoid cannabinoid CB(2) receptors in neuropathic pain. Elife, 9. 10.7554/eLife.55582

Carey, L. M., Xu, Z., Rajic, G., Makriyannis, A., Romero, J., Hillard, C., Mackie, K., & Hohmann, A. G. (2023). Peripheral sensory neuron CB2 cannabinoid receptors are necessary for both CB2-mediated antinociceptive efficacy and sparing of morphine tolerance in a mouse model of anti-retroviral toxic neuropathy. Pharmacological Research, 187. 10.1016/j.phrs.2022.106560

Cichon, J., Sun, L., & Yang, G. (2018, Mar 20). Spared Nerve Injury Model of Neuropathic Pain in Mice. Bio Protoc, 8(6). 10.21769/bioprotoc.2777

Dalm, B. D., Reddy, C. G., Howard, M. A., Kang, S., & Brennan, T. J. (2015, Dec). Conditioned place preference and spontaneous dorsal horn neuron activity in chronic constriction injury model in rats. Pain, 156(12), 2562–2571. 10.1097/j.pain.0000000000000365

Decosterd, I., & Woolf, C. J. (2000). Spared nerve injury: an animal model of persistent peripheral neuropathic pain. Pain, 87, 149–158.

Griggs, R. B., Bardo, M. T., & Taylor, B. K. (2015). Gabapentin alleviates affective pain after traumatic nerve injury. NeuroReport, 26(9), 522–527. 10.1097/wnr.0000000000000382

Guenther, K. G., Lin, X., Xu, Z., Makriyannis, A., Romero, J., Hillard, C. J., Mackie, K., & Hohmann, A. G. (2024). Cannabinoid CB2 receptors in primary sensory neurons are implicated in CB2 agonist-mediated suppression of paclitaxel-induced neuropathic nociception and sexually-dimorphic sparing of morphine tolerance. Biomedicine & Pharmacotherapy, 176. 10.1016/j.biopha.2024.116879

Guenther, K. G., Wirt, J. L., Oliva, I., Saberi, S. A., Crystal, J. D., & Hohmann, A. G. (2025). The cannabinoid CB2 agonist LY2828360 suppresses neuropathic pain behavior and attenuates morphine tolerance and conditioned place preference in rats. Neuropharmacology, 265. 10.1016/j.neuropharm.2024.110257

Guida, F., De Gregorio, D., Palazzo, E., Ricciardi, F., Boccella, S., Belardo, C., Iannotta, M., Infantino, R., Formato, F., Marabese, I., Luongo, L., de Novellis, V., & Maione, S. (2020). Behavioral, Biochemical and Electrophysiological Changes in Spared Nerve Injury Model of Neuropathic Pain. International Journal of Molecular Sciences, 21(9). 10.3390/ijms21093396

Guindon, J., & Hohmann, A. G. (2008, Jan). Cannabinoid CB2 receptors: a therapeutic target for the treatment of inflammatory and neuropathic pain. Br J Pharmacol, 153(2), 319–334. 10.1038/sj.bjp.0707531

Gutierrez, T., Crystal, J. D., Zvonok, A. M., Makriyannis, A., & Hohmann, A. G. (2011, Sep). Self-medication of a cannabinoid CB2 agonist in an animal model of neuropathic pain. Pain, 152(9), 1976–1987. 10.1016/j.pain.2011.03.038

Gutierrez, T., Oliva, I., Crystal, J. D., & Hohmann, A. G. (2021, Apr). Peripheral nerve injury promotes morphine-seeking behavior in rats during extinction. Exp Neurol, 338, 113601. 10.1016/j.expneurol.2021.113601

Kang, S. J., Kim, S., Lee, J., Kwak, C., Lee, K., Zhuo, M., & Kaang, B.-K. (2017). Inhibition of anterior cingulate cortex excitatory neuronal activity induces conditioned place preference in a mouse model of chronic inflammatory pain. The Korean Journal of Physiology & Pharmacology, 21(5). 10.4196/kjpp.2017.21.5.487

King, T., Vera-Portocarrero, L., Gutierrez, T., Vanderah, T. W., Dussor, G., Lai, J., Fields, H. L., & Porreca, F. (2009, Nov). Unmasking the tonic-aversive state in neuropathic pain. Nat Neurosci, 12(11), 1364–1366. 10.1038/nn.2407

Lin, X., Dhopeshwarkar, A. S., Huibregtse, M., Mackie, K., & Hohmann, A. G. (2018). Slowly Signaling G Protein–Biased CB2 Cannabinoid Receptor Agonist LY2828360 Suppresses Neuropathic Pain with Sustained Efficacy and Attenuates Morphine Tolerance and Dependence. Molecular Pharmacology, 93(2), 49–62. 10.1124/mol.117.109355

Lin, X., Xu, Z., Carey, L., Romero, J., Makriyannis, A., Hillard, C. J., Ruggiero, E., Dockum, M., Houk, G., Mackie, K., Albrecht, P. J., Rice, F. L., & Hohmann, A. G. (2022, May 1). A peripheral CB2 cannabinoid receptor mechanism suppresses chemotherapy-induced peripheral neuropathy: evidence from a CB2 reporter mouse. Pain, 163(5), 834–851. 10.1097/j.pain.0000000000002502

Martin, T. J., & Ewan, E. (2008, Oct). Chronic pain alters drug self-administration: implications for addiction and pain mechanisms. Exp Clin Psychopharmacol, 16(5), 357–366. 10.1037/a0013597

Navratilova, E., Xie, J. Y., King, T., & Porreca, F. (2013, Apr). Evaluation of reward from pain relief. Ann N Y Acad Sci, 1282, 1–11. 10.1111/nyas.12095

Park, H. J., Stokes, J. A., Pirie, E., Skahen, J., Shtaerman, Y., & Yaksh, T. L. (2013, Jan). Persistent hyperalgesia in the cisplatin-treated mouse as defined by threshold measures, the conditioned place preference paradigm, and changes in dorsal root ganglia activated transcription factor 3: the effects of gabapentin, ketorolac, and etanercept. Anesth Analg, 116(1), 224–231. 10.1213/ANE.0b013e31826e1007

Pereira, A., Chappell, A., Dethy, J., Hoeck, H., Arendt-Nielsen, L., Verfaille, S., Boulanger, B., Jullion, A., Johnson, M., & McNearney, T. (2013). A proof-of concept (poc) study including experimental pain models (epms) to assess the effects of a CB2 agonist (LY2828360) in the treatment of patients with osteoarthritic (oa) knee pain. Clin. Pharmacol. Ther. , 93, S56–S57. https://ascpt.onlinelibrary.wiley.com/doi/epdf/10.1038/clpt.2012.256

Rice, A. S. C., Finnerup, N. B., Kemp, H. I., Currie, G. L., & Baron, R. (2018). Sensory profiling in animal models of neuropathic pain: a call for back-translation. Pain, 159(5), 819–824. 10.1097/j.pain.0000000000001138

Wagner, K., Yang, J., Inceoglu, B., & Hammock, B. D. (2014). Soluble Epoxide Hydrolase Inhibition Is Antinociceptive in a Mouse Model of Diabetic Neuropathy. The Journal of Pain, 15(9), 907–914. 10.1016/j.jpain.2014.05.008

